# Single-nuclei RNA-seq on human retinal tissue provides improved transcriptome profiling

**DOI:** 10.1101/468207

**Authors:** Qingnan Liang, Rachayata Dharmat, Leah Owen, Akbar Shakoor, Yumei Li, Sangbae Kim, Albert Vitale, Ivana Kim, Denise Morgan, Nathaniel Wu, Margaret DeAngelis, Rui Chen

## Abstract

Gene expression profiling is an effective way to provide insights into cell function. However, for heterogeneous tissues, bulk RNA-Seq can only provide the average gene expression profile for all cells from the tissue, making the interpretation of the sequencing result challenging. Single-cell RNA-seq, on the other hand, generates transcriptomic profiles of individual cell and cell types, making it a powerful method to decode the heterogeneity in complex tissues.

The retina is a heterogeneous tissue composed of multiple cell types with distinct functions. Here we report the first single-nuclei RNA-seq transcriptomic study on human neural retinal tissue to identify transcriptome profile for individual cell types. Six retina samples from three healthy donors were profiled and RNA-seq data with high quality was obtained for 4730 single nuclei. All seven major cell types were observed from the dataset and signature genes for each cell type were identified by differential gene express analysis. The gene expression of the macular and peripheral retina was compared at the cell type level, showing significant improvement from previous bulk RNA-seq studies. Furthermore, our dataset showed improved power in prioritizing genes associated with human retinal diseases compared to both mouse single-cell RNA-seq and human bulk RNA-seq results. In conclusion, we demonstrated that feasibility of obtaining single cell transcriptome from human frozen tissues to provide additional insights that is missed by either the human bulk RNA-seq or the animal models.

## INTRODUCTION

Transcriptome profiling is a powerful tool for understanding gene function, classifying of cell type and state, and investigating human diseases^1–3^. Transcriptome can be more powerful when combined with other ‘omics’ data to build prediction models on human diseases^4^. For example, by combining transcriptomic data and proteomic data, a list of candidate disease genes can be predicted with high specificity^5^. However, until recently, vast majority transcriptome profiles are generated from profiling tissue samples containing thousands to millions of cells. Thus, gene expression information of individual cells would be lost. For tissues with high cellular heterogeneity, knowing the transcriptome profiles of each cell type would be important for both identification of novel cell types and understanding the functional organization of the tissue. Cell sorting would be required to obtain transcriptome of a single cell type; not only was it not always practical, but also the heterogeneities of many tissues were not fully revealed. This gap was met by the development of the high throughput single-cell RNA-seq technology^6–8^.

Transcriptomic studies on the single cell level was first performed decades ago^9, 10^, while the first single-cell transcriptome study based on Next-Generation Sequencing was reported ten years ago^11^. Since then, technologies have been dramatically improved in scale and sensitivity. Development in single-cell isolation, such as microfluidic-device-based methods, enables high throughput sequencing of thousands of cells at a time. Library construction methods, like SCRB-seq^12^ and SMART-seq^13^ allows for higher mRNA capture efficiency and lower bias. RNA-seq in single nuclei have been shown to be sufficient in representing the transcriptome of the whole cells and would facilitate transcriptome profiling when fresh samples, such as human samples, are not easily to obtain^14, 15^. Recently, single-cell transcriptome studies have the used in many applications, such as identify novel cell types^16^, reveal key players in cell differentiation^17^, and reconstruct the developmental trajectory in early embryonic development^18^.

The retina is an example of heterogeneous tissues and is composed of multiple neuronal and non-neuronal cell types^19^. In the human retina, there is an ordered array of approximately 70 different neuron types across 5 major classes: photoreceptor (rods and cones), retinal ganglion cells (RGCs), horizontal (HC), bipolar (BC), amacrine (AM) along with a non-neuronal Müller glial cell (MG), each playing a unique role in processing visual signal^19, 20^. Transcriptome of human retina has been reported using bulk tissue RNA-seq^21, 22^, and the overall gene expression profiles from different retinal regions (macular and peripheral region) were compared^23^. These studies provided the general transcriptomic information of the retina as a whole tissue, while they were not sufficient to reveal the complexity of the retina at individual cell type resolution. In addition, transcriptomic study of selected cell types of the human and primate retina were performed but were not sufficient to obtain the complete picture of all major cell types^24, 25^. However, the profiles of individual cell types, particularly types that account for a small portion of the retina, will offer important insights to the biology and disease. For example, it is often observed that one cell type, such as the cone cells in the cone-rod dystrophy (CRD) and the retinal ganglion cells in glaucoma, is the primary target of the disease^26, 27^. Given the great benefit of obtaining the transcriptome at individual cell type or even individual cell level, single-cell RNA-seq on human retina tissue is highly desired.

Recently, single-cell RNA-seq studies have been performed on mouse retina, both with the whole retina^28^ and with specifically enriched cell types^29, 30^. These studies provided unprecedentedly high-resolution transcriptomic data of each cell types and allowed for novel cell subtype discovery. The success of these studies supported the feasibility of single-cell studies on human retina using similar approaches. Despite the rich dataset from the mouse, it is essential to perform parallel study on the human retina given the considerable differences between the human and mouse. For example, the mouse retina lacks the macula region of the retina of humans and primates^31, 32^, a structure which is essential both for high visual acuity and color vision perception in the retina. Mouse cone cells are also different from human’s in their wavelength-sensitive opsin expression patterns. Here, we report the first transcriptomic study on healthy human retina tissues by using snRNA-seq. snRNA seq is an improvement over the standard single-cell RNA seq for profiling samples such as the neuronal tissue^33–36^. Using the snRNA-seq approach, a total of 4730 nuclei are profiled both the peripheral and macular region of three frozen human donor retina samples. Through unsupervised clustering of the gene expression profiles, clusters corresponding to all seven major cell types in human retina (rod, cone, MG, HC, AM, BC, and RGC) were identified. We compared the gene expression profile between macular and peripheral region, both as an entity and by individual cell type. While comparing the total macular and peripheral profiles, we found that the inherent differences in cell populations would strongly influence the detection of differentially expressed genes (DEG), a shortcoming of the bulk RNA-seq analysis. Single-nuclei study, on the other hand, allowed for macular-peripheral comparison within matching cell types. Significantly higher expression of mitochondrial electron transport genes was found in macular rod cells compared with peripheral ones, which might indicate that higher level of oxidation stress existed in the region and explain the vulnerability of macular rod cells as the stress accumulated as aging. In addition, as expected, compared to the published mouse single-cell data, the single-nuclei human data show stronger predictive power on genes associated with human disease. Finally, we found that photoreceptor DEGs significantly enriched inherited retinal disease (IRD) genes, indicating that it can serve as a prioritization tool for novel disease gene discovery and cell-specific pathway analysis. Overall, our study reported the first transcriptome profile of all major cell types of human retina at individual cell resolution, which would serve as a rich resource for the community.

## MATERIALS AND METHODS

### Macular and peripheral sample collection

As previously described in detail (Owen et al., in press), human donor eyes were obtained in collaboration with the Utah Lions Eye Bank. Only eyes within 6 hours postmortem time were used for this study. Both eyes of the donor undergo rigorous postmortem phenotyping including spectral domain optical coherence tomography (SD-OCT) and color fundus photography. Most importantly these images are taken in a manner consistent with the appearance of the analogous images utilized in the clinical setting. Dissections of donor eyes were carried out immediately according to a standardized protocol to reliably isolate the RPE/choroid from the retina and segregate the layers into quadrants^37^ (Owen et al., in press). After the eye is flowered and all imaging is complete, macula retina tissue is collected using an 6mm disposable biopsy punch (Integra, Cat # 33-37) centered over the fovea and flash frozen and stored at −80°C. The peripheral retina is collected in a similar manner from each of the four quadrants. To determine precise ocular phenotype relative to disease and or healthy aging, analysis of each set of images is performed by a team of retinal specialists and ophthalmologists at the University of Utah School of Medicine, Moran Eye Center and the Massachusetts Eye and Ear Infirmary Retina Service. Specifically, each donor eye is checked by independent review of the color fundus and OCT imaging; discrepancies are resolved by collaboration between a *minimum* of 3 specialists to ensure a robust and rigorous phenotypic analysis. This diagnosis is then compared to medical records and a standardized epidemiological questionnaire for the donor. For this study, both eyes for each donor were classified as AREDS 0/1 to be considered normal. Only one eye was used for each donor. Donors with any history of retinal degeneration, diabetes, macular degeneration, or drusen were not used for this study. Institutional approval for consent of patients to donate their eyes was obtained from the University of Utah and conformed to the tenets of the Declaration of Helsinki. All retinal tissues were de-identified in accordance with HIPPA Privacy Rules.

### Preparation of single-nucleus suspensions

Nuclei from frozen neural retinal tissue was isolated using RNase-free lysis buffer (10 mM Tris-HCl, 10 mM NaCl, 3 mM MgCl_2_, 0.1% NP40). The frozen tissue was resuspended in ice cold lysis buffer and triturated to break the tissue structure. The tissue aggregates were then homogenized using a Wheaton™ Dounce Tissue Grinder and centrifuged (500g) to pellet the nuclei. The pellet was re-suspended in fresh lysis buffer and homogenized to yield clean single-nuclei suspension. The collected nuclei were stained with DAPI (4’,6-diamidino-2-phenylindole, 10ug/ml) and were diluted to 1000 μl of 3E4/ml with 1× PBS (without Ca and Mg ions, pH 7.4, Thermo Fisher), RNase inhibitor (NEB, 40KU/ml) and Cell Diluent Buffer.

### The ICELL8™ single cell based single cell capture

Single nuclear capture and sequencing was performed on the ICELL8 single cell platform (Wafergen Biosytems). ICELL8 platform comprises of a multi-sample nano-dispenser that precisely dispensed 50nl of the single nuclei suspension into an ICELL8 nanowell microchip containing 5184 wells (150nl capacity). Assuming a Poisson distribution frequency for the number of cells per well, about 30% of the nanowells were expected to contain a single-nuclei under optimal conditions. Automated scanning fluorescent microscopy of the microchip was performed using Olympus BX43 fluorescent microscope with a robotic stage to visualize wells containing single nuclei (see Table 1 for single cell capture number across different experimental repeats). Automated well selection was performed using the CellSelect software (Wafergen Biosystems), which identified nanowells containing single nuclei and excluded wells with >1 nuclear, debris, nuclei clumps or empty wells. The candidate wells were manually evaluated for debris or clumps as an additional QC.

**Table 1.** Medical information of the sample donors.

| Donor Number | Globe | Age | Sex | Race |
| --- | --- | --- | --- | --- |
| 12-887 | OS (left eye) | 78 | F | Caucasian |
| 13-0025 | OD (right eye) | 78 | M | Caucasian |
| 13-0347 | OS (left eye) | 83 | M | Caucasian |

### Single-cell RT-PCR and library preparation

The chip was subjected to freeze-thaw in order to lyse the cells and 50 nl of reverse transcription and amplification solution (following ICELL8 protocol) was dispensed using the multi-sample nano-dispenser to candidate wells. Each well has a preprinted primer that contains an 11 nucleotide well-specific barcode which is added to the 3’ (A)n RNA tail of the nuclear transcripts during reverse transcription and cDNA amplification performed on candidate wells using SCRB-seq chemistry. After RT, the cDNA products from candidate wells were pooled, concentrated (Zymo Clean and Concentrator kit, Zymogen) and purified using 0.6x AMPure XP beads. A 3’ transcriptome enriched library was made by using Nextera XT Kit and 3’-specific P5 primer which was sequenced on the Illumina Hiseq2500.

### Data analysis

For bulk RNA-seq, FASTQ sequences were mapped to human genome hg19 (GRCh37) downloaded from UCSC genome browser website and aligned using STAR^38^. Transcript structure and abundance were estimated using Cufflinks^39–42^.

For snRNA-seq data analysis, FASTQ files were generated from Illumina base call files using bcl2fastq2 conversion software (v2.17). Sequence reads were aligned to the human genome hg19 (GRCh37). For transcriptome analysis, aligned reads were counted within exons using HTseq-count with default parameters^43^. All genes that were not detected in at least 2.5% of all our single cells were discarded for all further analyses. Cells were filtered based on a minimum number of 500 expressed genes per cell, a minimum number of 6000 and a maximum number of 100000 transcripts per cell. Data were log-transformed [log(read count + 1)] and normalized for all downstream analyses, using Seurat package (http://satijalab.org/seurat/)^44^. MultiCCA method in Seurat was used to align all the six datasets (3 macular, 3 peripheral). Aligned data were used for single cell clustering, tSNE visualization and differentially expressed gene (DEG) analysis. Without specific note, DEGs were assigned following three criteria: first, at least 25% of the group of interest expressed this gene; second, the gene in the group of interest should pass the Wilcoxon rank sum test against the reference group with Bonferroni adjusted p-value<0.05; third, the average fold-change of the gene in the group of interest against the reference group is more than 1.28 (logFC>0.25). Functional enrichment analysis was performed using DAVID^45^.

For mouse single-cell RNA-seq, the expression data was obtained from GSE63473^28^. Matrices from seven P14 mice, GSM1626793-1626799, were used for analysis. Gene filter of detection in >1% cell and cell filter of gene expression number over 500 were applied. CCA alignment and DEG analysis were performed with a similar procedure as the human sample.

## RESULTS

### Generation and the quality evaluation of the human retinal tissue single-nuclei RNA-seq

To generate transcriptome profiles for human photoreceptor cells, specially cones, retinae from three individual separate healthy donors were obtained (n = 3 donors). All three donors are Caucasian people and range from 60 to 80 years (Table 1) and were thoroughly examined as described in methods. Figure 1 shows an example of the donor tissues used for this study. There is no visible pathogenic indication, not even macular drusen, a hallmark of age-related macular degeneration, commonly found in this age group, according to the fundus images^46^. Two sample punches from each retina (n =6), one from the macula region and the other from the peripheral region, were collected and subjected to single cell nuclei RNA-Seq. After dispensing, nano-wells were imaged and only wells with single nuclei were selected. cDNA library construction and sequencing were performed for a total of 6,544 individual nuclei. Distribution of the number of nuclei from each sample is listed in Table 2. To exclude low quality data, several QC steps are conducted (detailed in methods). As results, 1,814 nuclei are filtered out, leaving a total number of 4,730 nuclei for downstream analysis. On average, 31,783 mapped reads are obtained per nucleus with the median number of genes detected at 1,054. To further evaluate the quality of snRNA-Seq data, bulk RNA-seq from the corresponding sample was performed. Good correlation between the bulk gene expression (FPKM, paired-end seq) and single-nuclei gene expression (average of normalized read count, 3’ end seq) was observed with positive correlation coefficient ranges from 0.57 to 0.72. This dataset, to the best of our knowledge, is the first snRNA-seq analysis on human retinal tissue.

**Figure 1.**
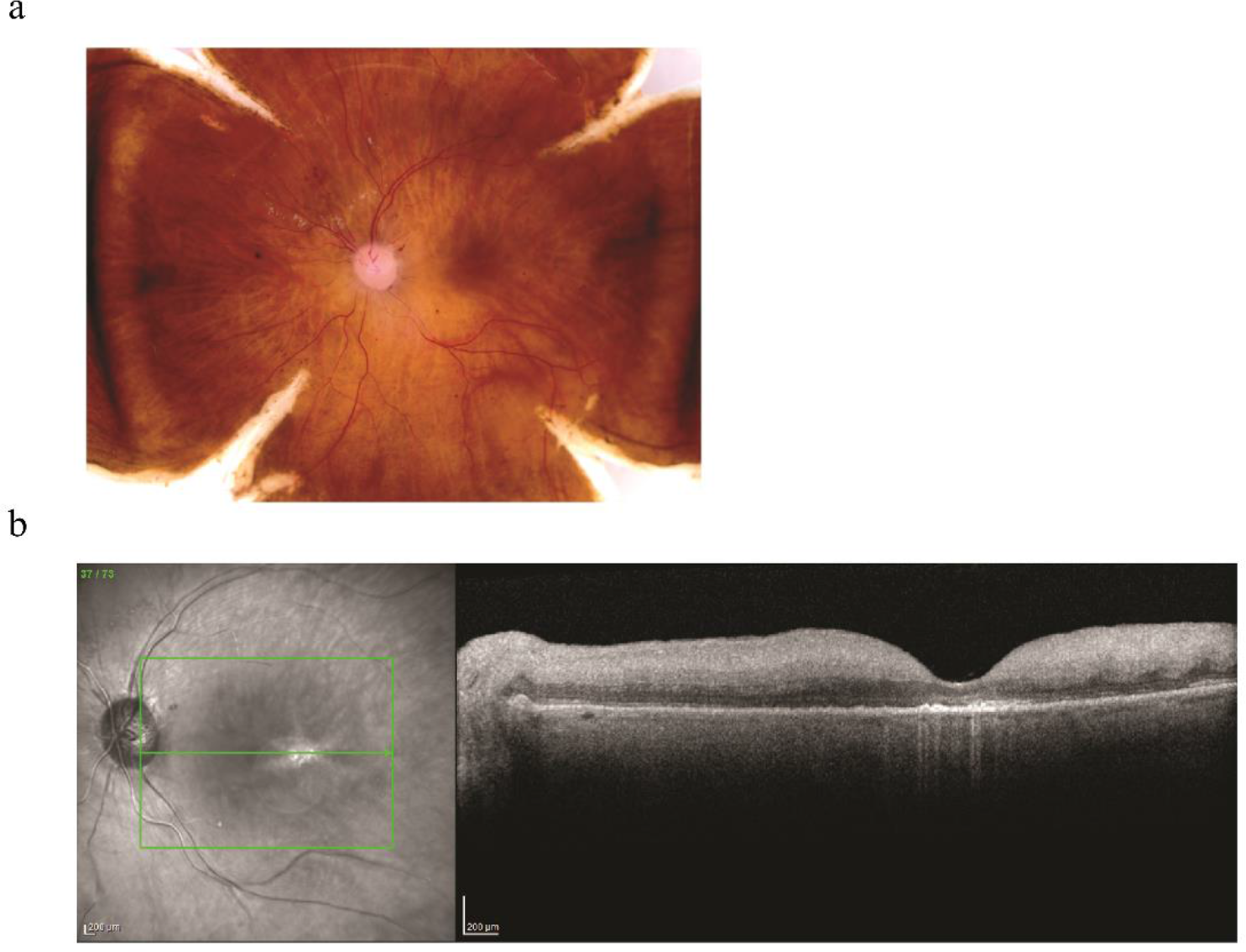
Representative analogous color fundus and OCT images demonstrating normal findings according to the Utah grading scale for postmortem eyes. A. Color fundus imaging of post mortem retina showing normal phenotype. B. OCT image of post mortem retina showing normal phenotype.

**Table 2.**
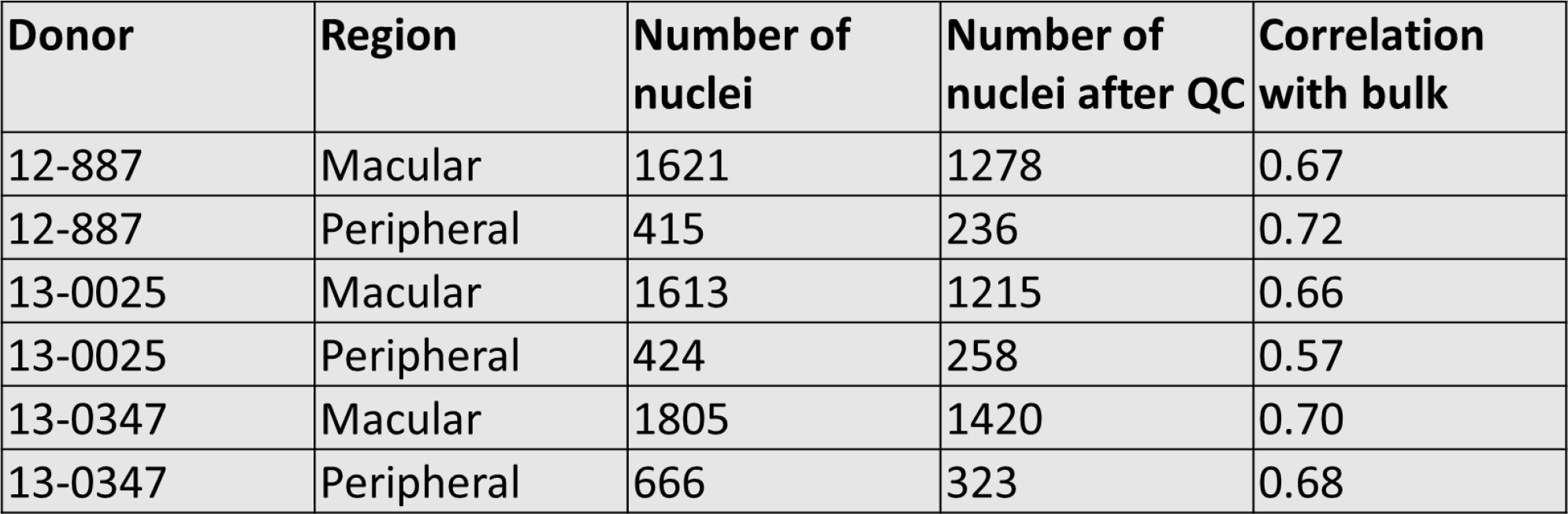
Basic sample information showing nuclei numbers and correlation with bulk RNA-seq of the same sample. Calculation of correlation was performed between the single nuclei RNA-seq and bulk RNA-seq of the same samples. For single nuclei RNA-seq (3’-end seq), the gene expression data from each nuclear were gene-filtered (2.5% cutoff, see method), normalized and log-transformed before averaging. For bulk RNA-seq (paired-end seq), FPKM was used.

### Unbiased single-cell transcriptomics profiling identifies all seven neuronal cell types of human retina

To identify individual retinal cell types, we performed unbiased clustering on the gene expression profiles of 4,730 human retinal nuclei. Seven clusters are identified, each of which contains cells from all six samples (Figure 2A), suggesting relatively low sample bias. Based on the expression pattern of cell type specific marker genes in the cluster, it is possible to map each cluster to individual cell types (Figure 2B). As shown in Figure 2C, markers for known retinal cell types, such as *PDE6A* for rod cells, *TRPM1* for bipolar cells (BC), *SLC1A3* for Müller glial cells (MG), *GAD1* for amacrine cells (AM), *SEPT4* for horizontal cells (HC), *ARR3* for cone cells and *RBPMS* for retinal ganglion cells (RGC), show cluster specific expression pattern. Thus, each cluster could be assigned to a known retinal cell type, which is further supported by examining additional known marker genes for each cell type (Supplement Table 1). Based on the number of nuclei in each cluster, we were able to quantify the proportion of each cell type in the sample. As shown in Figure 2D-E, composition of different cell types from human peripheral retina was generally consistent with that from previous mouse studies, with the exception of a higher percentage of MG cells and a lower percentage of AM cells being observed in the human retina^28, 47^. This trend is consistent with the results reported from a previous study in monkey, in which the relative ratio of BC: MG: AC: HC is close to 40:28:22:9^47, 48^. As expected, a lower rod proportion and higher BC, HC and RGC proportions are observed in the human macular sample compared with the human peripheral retina. A total of 122, 96, 126, 211, 228, 299 and 396 differentially expressed genes (DEGs) are identified for rod, BC, MG, AM, HC, cone, and RGC cell respectively (Supplement Table 2). Gene ontology enrichment analysis of biological process terms were performed with these DEGs (Supplement Table 3). Top GO terms enriched by each DEG lists showed consistency with our previous knowledge for each cell type, such as, for example, visual perception term for photoreceptor cells^49^, ion transmembrane transport term for retinal interneurons^50–52^, and MAPK pathway regulation term for Müller glia cells^53^. These results all support the data quality is sufficient to faithfully represent the transcriptome profiles of major cell types of human retina.

**Figure 2.**
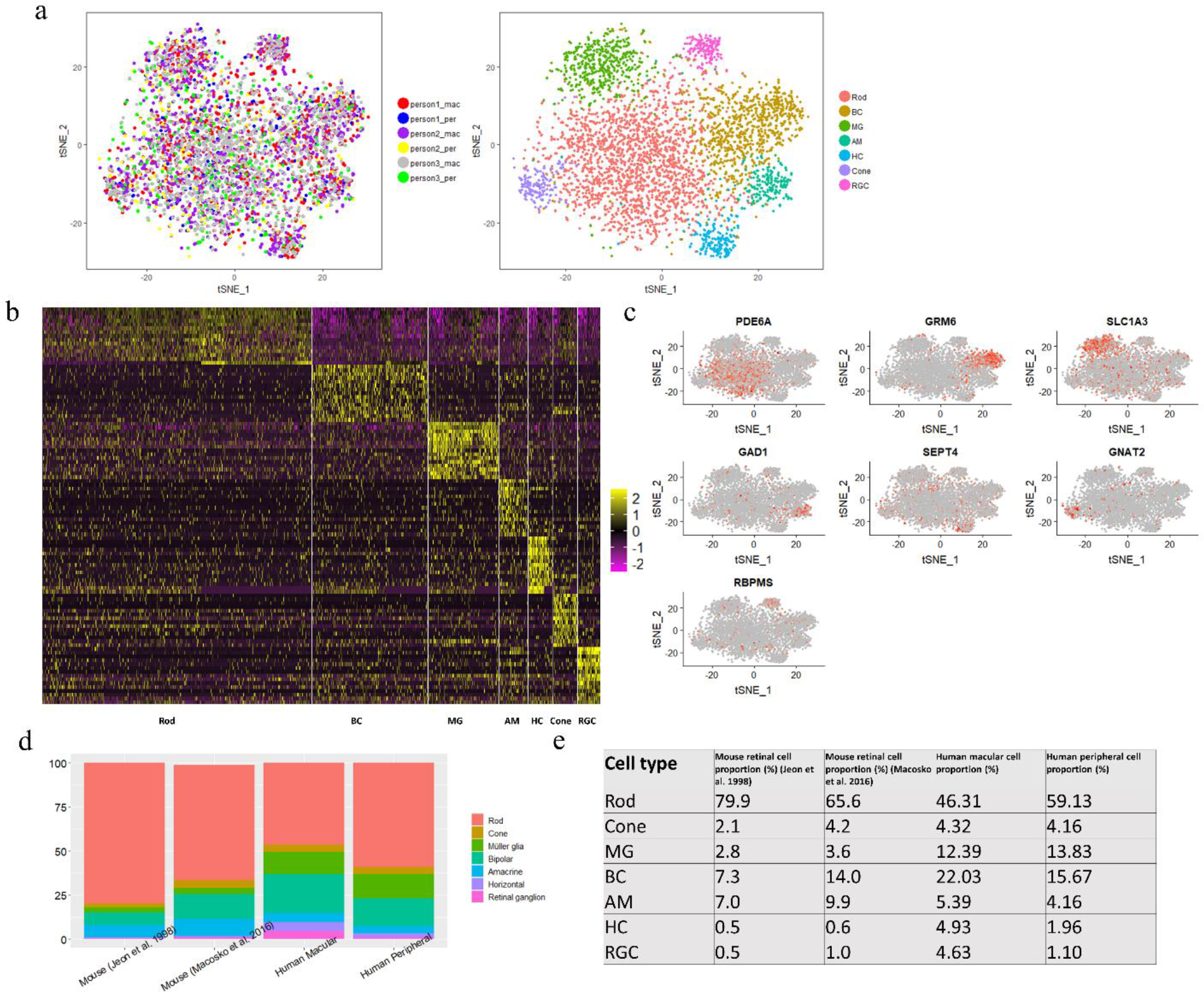
Unsupervised clustering identifies 7 major cell types in human retina. A. Clustering of 4730 human retina single nuclei expression profiles into 7 populations (left) and representation of alignment of 6 dataset from 3 donors. B. Heatmap of top 15 DEGs in each cell type. Each column represents a cell while each row represents a gene. Gene expression values are scaled. C. Profiles of known markers (*PDE6A, GRM6, SLC1A3, GAD1, SEPT4, ARR3, RBPMS*) in each cluster. D-E. Comparison of proportion of 7 major cell types of mouse and human retina.

### Differentially expressed genes in the macular region compared with peripheral region

The macula is a structure unique to human and other primates. Several studies aiming at identifying genes that are differentially expressed between macular and peripheral retina have been conducted. For example, using a SAGE approach, Sharon et al reported 20 genes with high expression in the macula and 23 genes with high expression in peripheral region in the retina (referred as SAGE gene list)^54^. In addition, based on bulk RNA-Seq, Li et al reported 1,239 genes with high expression in macular retina and 812 genes with high expression in the peripheral macular region^23^. To evaluate our data to these published data, we generated the virtual bulk macular and peripheral RNA-Seq data by in silico combining snRNA-Seq from each region. As a result, we obtained a list of 181 genes that are highly expressed in the macular region and 118 genes that are highly expressed in the peripheral region (to get the most significant DEGs, we only included genes with 1.5-fold changes and adjusted p-value<0.01, Supplement Table 6). Significant overlapping between the SAGE gene list and our list is observed. Specifically, for the SAGE gene list, 11 out of 20 macular genes (*CPLX1, D4S234E, NEFL, STMN2, UCHL1, NDRG4, TUBA1B, DPYSL2, APP, YWHAH, MDH1*) and 11 out of 23 peripheral genes (*SAG, RCVRN, UNC119, GPX3, PDE6G, ROM1, ABCA4, DDC, PDE6B, GNB1, NRL*) are also observed in our gene list. Based on GO term analysis, the macular gene list is enriched in the cholesterol biosynthetic process, microtubule skeleton organization, microtubule-based movement (Figure 3A), indicating a more interneuron-like transcriptome, while the peripheral genes are enriched for visual perception, phototransduction and rhodopsin related signal transduction pathways. Similar GO terms, such as ion transport in macular genes and visual perception in peripheral genes (Figure 3A), are enriched in both our data set and the Li dataset (gene set not public available).

**Figure 3.**
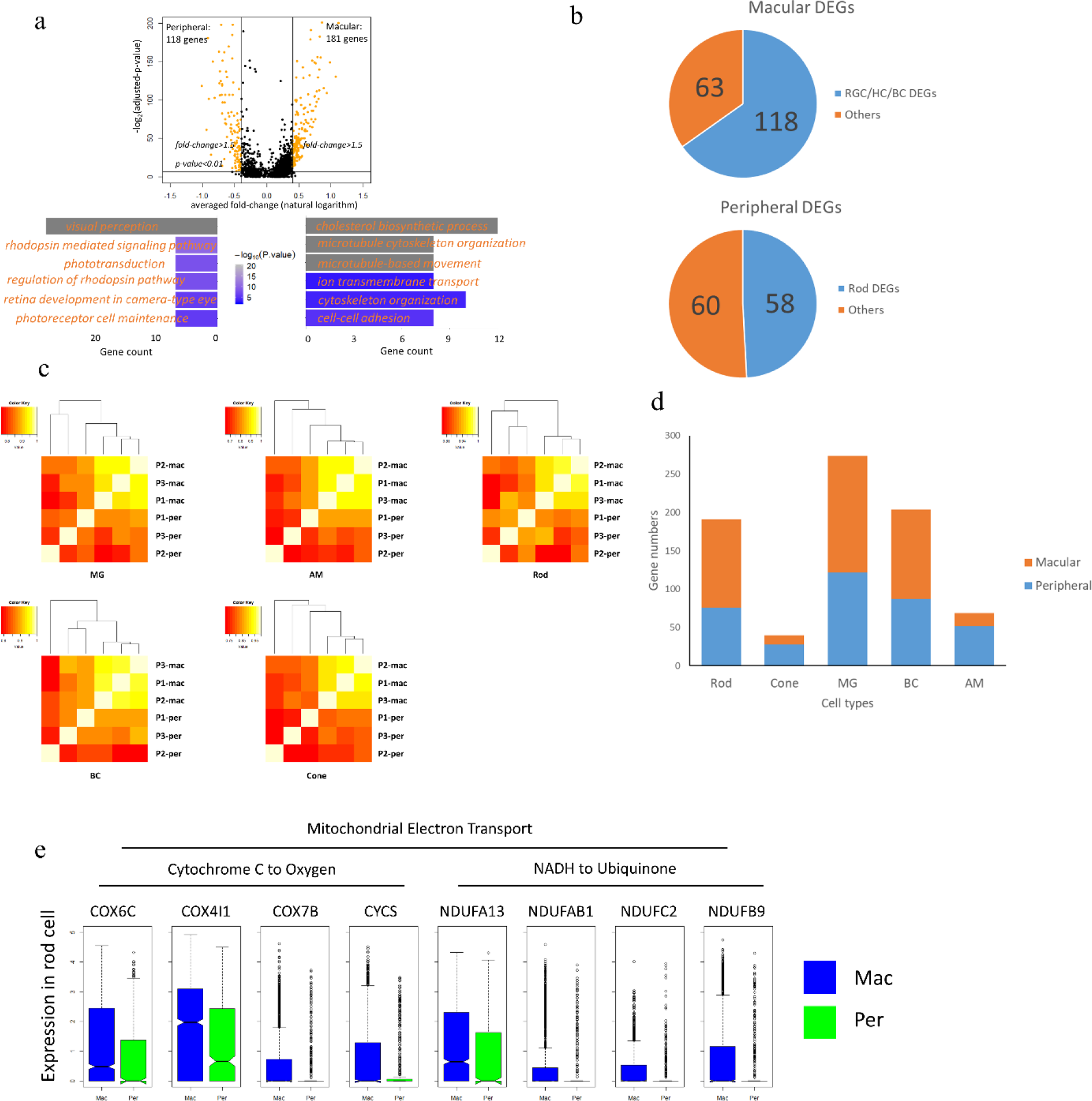
Transcriptome difference revealed between macular and peripheral region. A. Differential expressed genes between the overall macular and peripheral cells and biological process GO term enrichment. B. Proportion of cell type DEGs in the overall macular-peripheral DEGs showing cell population affection on DEG detection. C. Heatmap demonstrating correlation of averaged gene expression of five major cell types in six samples. D. Number of cell type specific DEGs in rod, cone, amacrine cell (AM), bipolar cell (BC) and Müller glia cell (MG). E. Demonstration of mitochondrial electron transport related genes showing differential expression in rod cells in macular and peripheral region.

Based on previous report and our data, although the macular and the peripheral region have the same types of cell, the proportion of cell types are different between macular and peripheral human retina (Figure 2D-E). As a result, genes that show differential expressing level between the macular and peripheral regions in bulk transcriptome profiling experiments might be due to difference in cell proportion instead of true expression level difference. In the 181 macular DEGs, 118 were also found in DEG lists of RGC, BC or HC, which are much more abundant in the macular region (Figure 2D-E), while in the 118 peripheral DEGs, 58 are found as rod DEGs (Figure 3B). This result confirmed that “bulk” RNA-seq data tended to be affected by cell population variations in different regions in the retina and indicated that the single-cell level study is required for revealing genuine macular-peripheral similarities and differences. Our snRNA-Seq data overcomes the issue and allows direct comparison of gene expression between macular and peripheral retina for the same cell type. Correlation of averaged gene expression patterns of five cell types, rod, cone, MG, BC and AM, from each sample were calculated (Figure 3C). The profiles of cells from the macular and peripheral form two sub-clusters, indicating differences do exist within the same cell type depending on the region. Within each cell types (rod, cone, MG, BC and AM) between the macular cells and peripheral cells, we performed differential expression analysis and found cell type DEGs (Figure 3D, Supplement Table 6). Among the DEGs, interestingly, some mitochondrial electron transport related genes showed high expression in macular rod cells compared with peripheral ones (Figure 3E). Rod photoreceptor cells have large numbers of mitochondria packed in the inner segment^55^ for they require high amount of energy in order to keep the high turnover rate of the outer segment and support phototransduction^56^. Higher expression level of these genes indicates higher oxidative stress which has been linked to photoreceptor death^57^. Mitochondria is the major source of oxidative stress in the retina and it could get accumulated as the organism ages^58^. Mitochondrial dysfunction in retinal pigment epithelial cells (RPEs) has been associated with retinal diseases like age-related macular degeneration (AMD), while the effect in photoreceptor cells remains elusive^56^. Our finding might be able to contribute to the explanation of why macula photoreceptors are more vulnerable then peripheral ones. In conclusion, the cell-type based macular-peripheral comparison reveals regional variances that bulk RNA-seq cannot achieve.

### DEG analysis reveals human cone-rod differences and shows increased relevance to human disease

More than half of the retina cell population is composed of photoreceptor cells. By combining rod and cone photoreceptor cells, a list of 147 genes that are highly enriched in photoreceptors was obtained (Supplement Table 2). These genes show significant enrichment of biological process GO terms of visual perception, rhodopsin mediated signaling pathway, maintenance of photoreceptor cells and regulation of rhodopsin mediated signaling pathway (Figure 4A). This is largely consistent with the known function of photoreceptor cells, which further validate our cluster assignment and DEG analysis.

**Figure 4.**
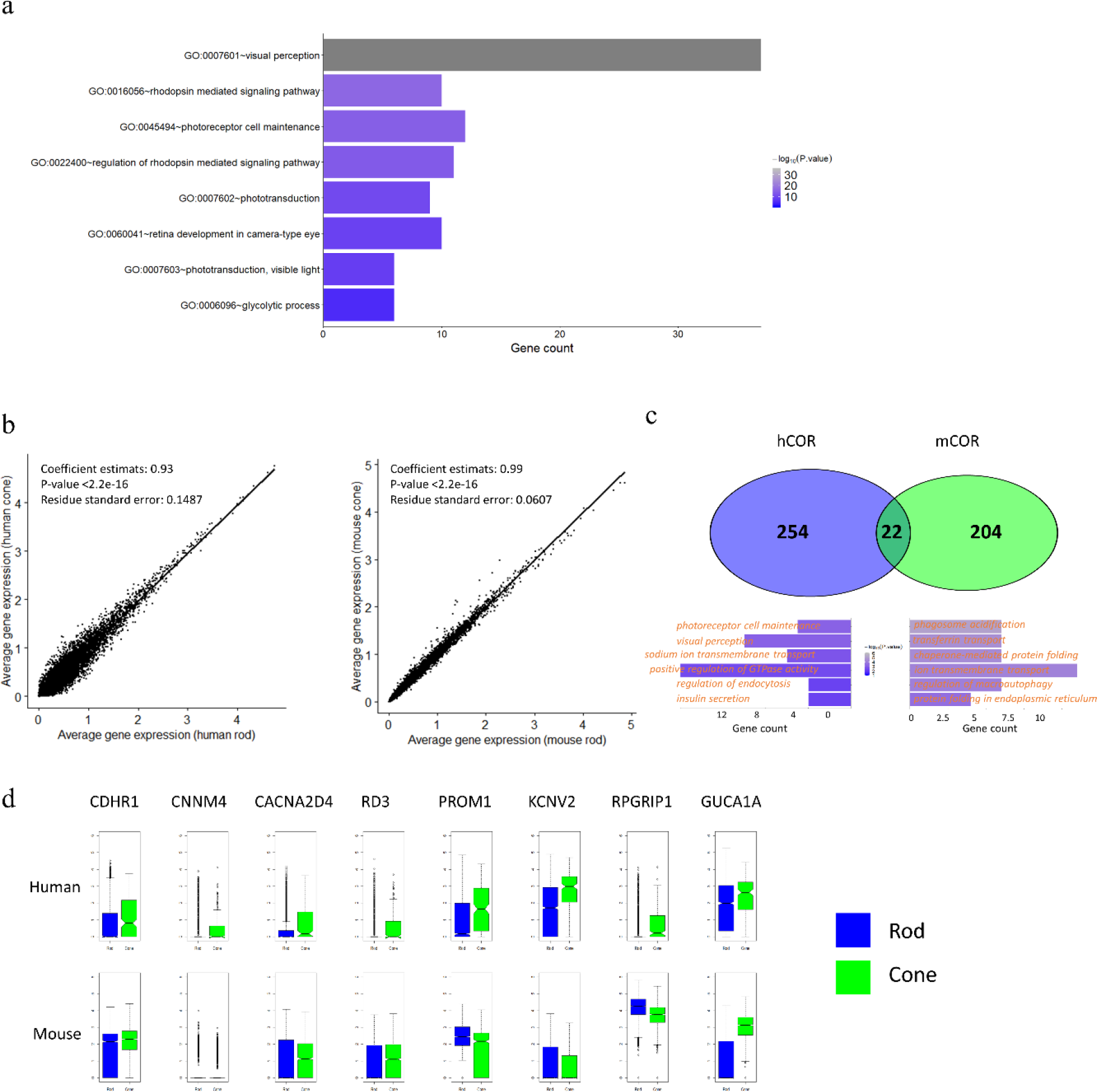
Differentially expressed genes are revealed in rod and cone photoreceptors. A. Biological process GO term enrichment by photoreceptor cell DEGs. B. Gene expression correlation between rod and cone cells in human and mouse. C. Overlap of the hCOR (human cone-over-rod) and mCOR (mouse cone-over-rod) gene list and biological process GO term enriched in non-overlapping part. D. Demonstration of the expression of CRD/LCA genes in the non-overlapping part of hCOR (*CDHR1, CNNM4, CACNA2D4, RD3, PROM1, KCNV2, RPGRIP1*) and mCOR (*GUCA1A*).

In photoreceptor cells, cones are functionally important^19, 49, 59, 60^ and less studied. During the retina development, photoreceptor precursors first commit to the photoreceptor cell fate, and then get specified to subtypes, rods and cones^61^. Thus, cone cells and rod cells are close in development and share similarities in functions, and comparison of transcriptomes between cone cells and rod cells would provide informative insights to cone cell specific functions. With published single-cell RNA-seq data of mouse retina^28^, rod and cone cells were identified with similar methods as that we use for the human data. Thus, we first compared the average gene expression between human rod and cones, and between mouse rod and cones (Figure 4B). High Spearman correlations have been found between each pair, indicating the overall transcriptome similarity between rod and cone cells. In the linear models that using rod (or cone) gene expression as the observation value and cone (or rod) gene expression as the only variable, the human data showed a residue standard error (0.1487) 2.4 times as much as the mouse data (0.0607), meaning that the gene expression values between human rod and cone are more disperse. In other words, in comparison, human cones are more distinct from human rods, while mouse rod and cones are more similar.

Genes with higher cone expression compared with rod may be relevant to cone specific functions. We identified 276 genes that are highly expressed in human cone cells compared with rod cells (human cone-over-rod gene list, referred as hCOR list, Supplement Table 5). GO analysis on biological process terms revealed that these genes enrich terms such as visual perception, response to stimulus, sodium ion membrane transport, and cilium assembly (Supplement Table 4). Previous study in mice retina has identified 226 genes that are be highly expressed in cone cells compared with rod cells (mouse cone-over-rod list, referred as mCOR list, Supplement Table 5). Interestingly, this mCOR list only share about 10% with the hCOR list (Figure 4C), indicating the considerable differences between human and mouse cone cells. Excluding the 22 shared genes by hCOR and mCOR, the rest genes in hCOR enrich biological process GO terms such as photoreceptor cell maintenance, visual perception and sodium ion transport, while the top terms enriched by the rest of mCOR list were not as phototransduction-related. This result again indicates that in the transcriptome level, human cone and rod cells are more distinct, while mouse cone and rod cells are closer. Besides, in the hCOR list (excluding shared genes), 7 genes (*CDHR1, CNNM4, CACNA2D4, RD3, PROM1, KCNV2, RPGRIP1*) are found as known human CRD or LCA genes, while mCOR list only has one (*GUCA1A*). As an example, high expression of *RPGRIP1* and *RD3*, both being known human IRD genes^62–71^, are showing significantly higher expression in human cones compared to rods, as is shown in Figure 4D. Consistent with the expression pattern, patients with mutations in *RPGRIP1* and *RD3* show LCA and CRD phenotype, which has more severe defects in cones than rods. In contrast, in the mice dataset, these two gene show no differential expression in rod and cone (Figure 4D). Consistently, KO mouse models of these two genes are reported to display RP like phenotypes^71–73^, which were resulting from rod cells early affections. Therefore, the differences in mouse and human phenotype is likely due to difference in cell specific expression of the gene. Thus, the human cone profile we report here show increased relevance to human diseases and would serve as a better resource for human study.

### Differentially expressed genes in photoreceptor cells enrich inherited retinal disease genes

With the expression profile for each retinal neuronal cell type generated in this study, we sought to examine its potential utility in identifying genes associated with human retina diseases. A gene list of 234 genes that contains all the known IRD genes for RP, LCA, CRD, and other retinopathies was obtained from the retnet (RetNet, http://www.sph.uth.tmc.edu/RetNet/) (Supplement Table 7). As expected, robust expression of most of these known IRD genes (197 of the 234) could be detected in our dataset (Supplement Table 7). Additionally, the detected IRD genes are expressed at significantly higher level than average (Figure 5A, p-value = 2.27e-07, two-sample t-test). Since the vast majority of known IRD associated genes affects photoreceptor cells exclusively, we reason that significant overlap should be detected between IRD genes and the photoreceptor enriched gene set. Indeed, 44 IRD genes are among the 147 PR DEGs, representing a significant enrichment over background (odds ratio = 29.62, p-value<2.2e-16, fisher’s exact test) (Figure 5B, Supplement Table 8). Thus, we propose that the PR DEG list could also be directly used as a gene prioritization list for novel IRD gene discovery. This result showed improvement compared with previous bulk RNA-seq studies. Pinelli et al performed RNA-seq on 50 retina samples and used co-expression analysis to predict potential IRDs and were able to recover 56 known genes from a list of 472 genes, with an odds ratio of 15 (Figure 5B)^22^. In comparison, our PR DEG list, without any optimization, show highly enrichment level with the odds ratio almost doubled (z score = 2.54, p-value = 0.0055).

**Figure 5.**
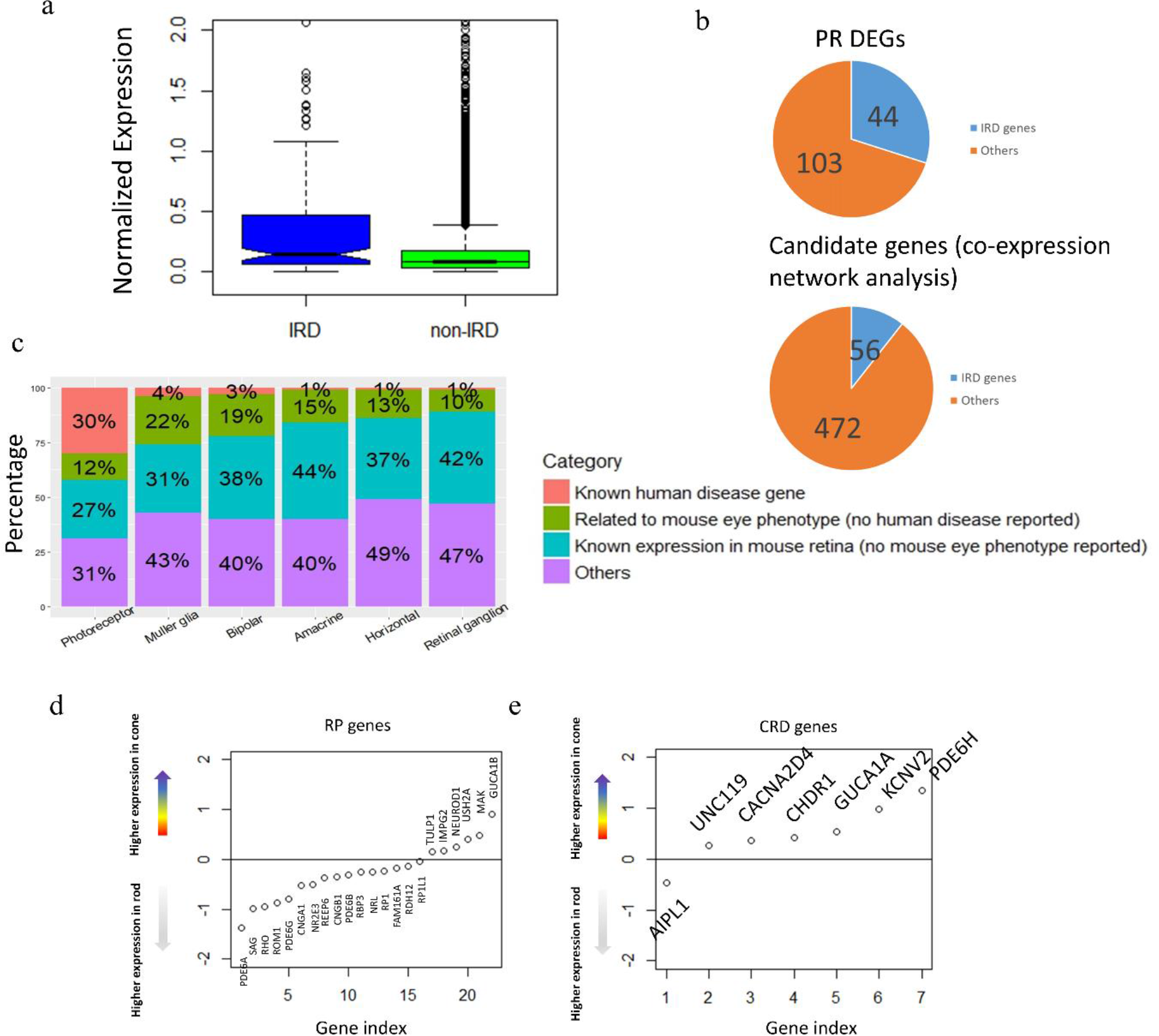
Photoreceptor DEGs enrich human IRDs. A. IRD genes generally show higher expression level than the rest of genes in human retina. B. Photoreceptor cell DEG list enrich known IRD genes and showed improvement from previous study based on bulk RNA-seq C. Distribution of human retinal disease genes, mouse eye phenotype genes (but no human disease discovered), mouse retinal expressing genes (but no eye phenotype found) in DEGs of all cell types (PR is merged from rod and cone). D. RP genes and CRD genes show different expression trend in rod and cone photoreceptors. The y-axis is representing the differential expression of each gene in cone cells compared with rod cells (cone minus rod). Expression values were log-transformed followed by averaging within cell types (rod, cone).

Depending on the timing and severity of rod and cone photoreceptors defect, IRDs can be classified into different subtypes clinically. For example, although cone photoreceptor cell degeneration is observed as the disease progresses, RP is primarily due to defect in rod photoreceptors^74–76^. On the other hand, cone-rod dystrophy (CRD) is result from cone degeneration from the very beginning^26, 27^. Among the 44 IRD genes that show photoreceptor cell-specific expression, 26 have been associated with RP while 11 have been associated with CRD. Consistent with the clinical phenotype, on average, RP associated genes have higher expression in rod cells while CRD associated genes show higher in cone cells (Figure 5C-D).

Given the significant enrichment of IRD associated genes in the cell type DEG set, it can be potentially used to prioritize candidate IRD disease genes. To further test this idea, the eye phenotype for mice with photoreceptor cell DEG orthologous gene mutation is examined. For the 103 photoreceptor DEGs that have not been associated with human retina diseases, 49 have knock out mice recorded in the MGI database with non-lethal phenotypes. Among them, 18 shown phenotypes in the visual system (Supplement Table 8), such as *GNGT1*, *RDH8* and *RCVRN*. Protein encoded by *GNGT1* is known to locate in the outer segment of photoreceptor cells and plays important role in phototransduction^5, 77^. *RDH8* gene encodes a human retinol dehydrogenase, and mutation of its mouse ortholog causes delayed rod function recovery after light exposure and accumulation of all-trans-retinal in the rod outer segment^78^. *RCVRN* is a calcium-binding protein which is a regulator of rod sensitivity of dim lights, and it is also found to be an auto-antigen of cancer-associated retinopathy^79^. In addition to investigating the genes with mouse eye phenotype, we also highlighted the genes with known mouse retina expression (MGI database query). In summary, our photoreceptor cell DEG list would serve as a prioritization tool for novel disease gene discovery.

### Retinal disease genes and mouse phenotype related genes in other retinal cell types

In contrast to the significant overlap between IRD genes and photoreceptor (PR) DEGs, no significant enrichment is observed in DEGs for other retina cell types. Although rare, it has been shown that defects in cell types in the neural retina other than PR can also lead to IRD. Indeed, 14 known IRD genes show enriched expression in cell types not restricted to photoreceptor cells (Supplement Table 8). From those genes, we found a few of them have known functions consistent with the expression pattern and human phenotype. For example, *RDH11* is highly expressed in Müller Glia cell which plays an important function in retinal cycle and mutations in *RDH11* lead to RP^80^. *GRM6* and *TRPM1* are bipolar DEGs and their mutations could cause a recessive congenital stationary night blindness (CSNB), and CSNB is known to be caused by abnormal photoreceptor-bipolar signaling^81–87^. Besides, in their DEG lists, 28, 30, 38, 32, and 18 genes that are highly expressed in MG, HC, RGC, AM and BC, respectively, showed correlation to the mouse eye phenotypes (Supplement Table 8). These genes could potentially serve as prioritization of possibly disease-causing genes for further study of non-PR-related IRDs.

## DISCUSSION

Studying the cell type specific transcriptome expands our understanding of the cell function within heterogeneous tissues, which the retina is a good example for. The retina contains seven major cell types with various proportion: rod cells consists of over half while some types such as amacrine cells and RGCs are found to be much less. All the cell types have distinct functions and coordinate to allow for visual perception and regulation. In multiple pathological conditions, especially inherited retinal diseases (IRDs), certain cell types, such as rod, cone, or bipolar cells, but not the whole tissue, are affected at the early stage^26, 74, 88^. Thus, understanding the transcriptome will also facilitate disease-related studies. In the retina, transcriptome profiles of many cell types, especially rare ones, are usually masked by the average of all cell populations in bulk RNA-seq of the whole retinal tissue. Cell-surface-marker-based sorting and purification methods have been developed for enriching for all cell types. Thus, single-cell RNA-seq stands out as the most optimal and unbiased method for obtaining the transcriptome of the cell types in the retina.

We report the first transcriptome profiles of the human retinal major cell types at the single-cell level from 4,730 nuclei. These nuclei were from six samples originating from three donor retinae. It is worth emphasizing that the tissues we used all came from similar genetic background, all underwent post-mortem phenotyping, and are shown to be normal. As standard in current clinical practices, OCT images are used to resolve the appearance of subretinal drusen, fluid, atrophy and fibrosis and differentiate artifact from pathology. These images (Figure 1) demonstrate the usage of a combination of fundus and OCT imaging techniques for post-mortem eye common in clinical practice according to the Utah Grading Scale for Post mortem Eyes (Owen et al In press IOVS). All major cell types were found in every sample with the reasonable proportion and expected marker expression. Thus, the transcriptome profiles we demonstrated here are reproducible and reliable.

Regional transcriptomes for tissues are of great interest for researchers. Here, we demonstrated that for heterogeneous tissue, single-cell studies out-perform bulk studies in regional specific transcriptome profiling. Bulk studies are largely limited by variations of cell population in different regions so that the genes showing variable expression levels could just be the outcome of changed cell proportions. Single-cell studies, on the other hand, allows for comparisons between certain cell types and facilitates detection of real differences.

The transcriptome profiles of cell types in the human retina are also major findings in our study. Mice, the most studied animal model for retinal degeneration, are not an ideal model for studying cone biology as its cones have significant differences from those in humans. For example, in the human retina, cone cells are highly enriched in a region called the fovea near the central macula which is absent in the mouse retina^31, 89^. In addition, mice have two types of cone-opsins, namely *Opn1sw* for short-wavelength sensing and *Opn1mw* for middle- and long-wavelength sensing. Some mouse cone cells express only one type of opsin while a considerable proportion (40% for C57/BL6 mice) express both^90^. In contrast, humans have three types of cone-opsins, *OPN1SW*, *OPN1MW* and *OPN1LW*, and each cone cell only express one type of opsin^61^. Mustafi et al previously compared the transcriptomes between cone-enriched macular region and rod-enriched peripheral region in monkey to analyze rod and cone signatures^25^. They found known cone makers, such as *SLC24A2* and *OPN1MW*. However, they also found other genes, including *NEFL* and *NEFM*, which generally showed higher expression levels in the macular region and might partially be driven by the uneven proportion of RGCs. Welby et al developed a sorting method to enrich fetal cones in human and reported the transcriptome of these fetal cone cells^24^. Their findings identified the cone signature during the development (9 to 20 weeks post conception), which could be useful in recapitulating some aspects of the human adult cone. Our study took advantage of “in silico” sorting to separate cones from other cell types and thus provided better profiling.

The utility of snRNA-seq on human retinal tissue is assessed in our study. snRNA-seq is a viable alternative method to single-cell RNA-seq which is more practical for human tissue study because it could be applied to frozen neuronal tissue. In addition, snRNA-seq has less bias on sampling as it is not affected by factors such as cell size. Gao et al reported that, in breast cancer cells, the snRNA-seq was representative of single-cell RNA-seq ^15^. Our single-nuclei profiles showed a good correlation with bulk RNA-seq of the same sample, and the cell type profiles also displayed consistency with published human retinal cell markers (Supplement Table 1).

The transcriptome profiles of human retinal cell types could be a useful tool for researchers. For example, we observed significant enrichment of IRD genes among the genes with specificity in photoreceptor cells. In addition, the rest of the list also contains multiple genes that were linked to mouse eye phenotypes. Thus, we would use this list as a reference in prioritizing genes for novel IRD gene discovery. Additionally, it should not be neglected that some genes with specificity in other cell types were also found to be IRD genes or cause mouse eye phenotype. Besides the uses we addressed here, our findings could also be useful for other studies. For instance, the cell type markers reported in our data would facilitate approaches in cell purification of the human retina, which may lead to better cell type profiles in the future.

## Author Contributions

RC and MD conceived the project. QL, SK, RD and NW did the data analysis. LO, AS, AV, IK, DM and MD collected the human donor retinas, performed the phenotyping and dissection. RD and YL prepared the nuclei sample and performed the single-nuclei RNA-seq. QL, RD, MD and RC wrote the manuscript with input from all other authors. All authors proofread the manuscript.

## Acknowledgement

This work is supported by grants from Retina Research Foundation (R.C.) and NEI (R01EY018571 and R01EY022356 to R.C.). Bulk and single-cell RNA-Seq were performed at the Single Cell Genomics Core at BCM partially supported by NIH shared instrument grants (S10OD018033, S10OD023469) and P30EY002520 to R.C.

